# A High-Throughput Automated Pipeline to Analyze Synapse Function by Calcium Imaging

**DOI:** 10.64898/2026.03.16.712134

**Authors:** John Carl Begley, Harald Prüss, Paul Turko, Camin Dean

## Abstract

Synapses are the basic unit of information transfer between neurons. Their dysfunction is a common trigger of cognitive diseases and disorders. However, high-throughput analysis methods to assess synaptic function and dysfunction are lacking. Calcium imaging in cultured neurons in the absence of Mg^2+^ and presence of TTX allows visualization of NMDAR-dependent spontaneous synaptic calcium transients, which report pre and postsynaptic function. Here, we introduce a high-throughput automated analysis pipeline that combines Suite2p ROI detection and Python scripts to analyze tens of thousands of synapses and quantify changes in presynaptic vesicle fusion rates (frequency), postsynaptic function (amplitude), and the number of functional synapses. We use this pipeline to test known NMDAR agonists (glycine) and antagonists (ketamine, memantine, APV), presynaptic function modulating compounds (PDBu), and encephalitis patient-derived NMDAR auto-antibodies, where our pipeline proved more sensitive in detecting dysfunction at the single-synapse level than other methods. The ability to detect, track, and quantify activity across tens of thousands of synapses and millions of synaptic calcium transients using this pipeline will aid drug discovery of compounds that protect synapse function.

## Introduction

Calcium imaging is a well-established method for studying neuronal activity and connectivity. It is routinely used to infer electrical activity across large populations of neurons in vivo and in vitro in place of, or alongside, more invasive and low-throughput electrophysiological recordings. The improved sensitivity of genetically-encoded calcium indicators allows the detection of subthreshold changes in intracellular calcium concentrations in neurons and their processes (Li et al., 2019). Synaptic calcium imaging can be achieved by imaging in a modified artificial cerebrospinal fluid (ACSF) solution that contains tetrodotoxin (TTX), to block action potentials, and 0 mM Mg^2+^, to remove the magnesium block of NMDA receptors (NMDARs). This allows NMDARs in dendritic spines to report spontaneous glutamate release from presynaptic vesicles by calcium influx into postsynaptic dendrites.

The combination of 0 Mg^2+^ and TTX was initially used in electrophysiological experiments to demonstrate that NMDARs are the primary glutamatergic receptors to flux calcium (Iino et al., 1990; Mayer et al., 1984). Removing magnesium to ungate NMDARs was then co-opted for calcium imaging to study synaptic calcium transients in vivo in response to glutamate uncaging (Sobczyk et al., 2005). This method was later adapted to visualize synaptic calcium transients in vitro (Andreae & Burrone, 2015). Synaptic calcium imaging has been applied in vitro to investigate NMDAR trafficking and activity in developing dendrites (Andreae & Burrone, 2015), the effect of beta amyloid (Prikhodko et al., 2024; Sinnen et al., 2016), and the effect of a patient-derived NMDAR autoantibody (Dean et al., 2022) on synaptic calcium transients, for example.

Despite the adoption of this method by multiple research groups, the analysis of synaptic calcium imaging recordings has remained low-throughput. To the best of our knowledge, all previous uses of this method have relied on either manual or semi-manual approaches to identify potential synaptic regions of interest (ROIs) in combination with custom-written code and software for analysis. These manual methods are not only time- and labor-intensive but also susceptible to user bias and may potentially miss small, nuanced changes in synaptic activity due to small sample sizes. Manual identification of ROIs is also incompatible with larger datasets, leaving previous studies to rely on, at most, datasets of 1000-2000 synapses per experimental condition. Most neurons form thousands of synapses, so there is clear room for optimization. Without the limitations of manual detection and low-throughput analysis, this method could be used for large-scale screenings of compounds and their effects on synapses by reporting functional synapse number, presynaptic vesicle fusion rates and neurotransmitter release properties, and postsynaptic receptor function.

We therefore sought to automate the detection and analysis of synaptic calcium events with a single open-source pipeline that would allow all visible synapses to be detected and analyzed with minimal user intervention. This allowed us to curate datasets with tens of thousands of individual synapses. From these datasets, we could investigate changes to individual synapses and populations of synapses before, during, or after treatment with potential synapse-modulating compounds. We adapted the open-source Python-based somatic calcium imaging analysis program Suite2p (Pachitariu et al., 2016) to detect synaptic ROIs. Then, we developed and validated an automated workflow in Python combining Suite2p ROI detection and analysis of spontaneous synaptic calcium transients in neuronal cultures expressing GCaMP6f. Using these scripts, we were able to identify synaptic calcium transient sites and calculate frequency and amplitude values for calcium transients at individual synapses. Moreover, we also report changes in active synapse numbers.

With this analysis pipeline, we reliably report NMDAR-dependent synaptic calcium transients, benchmarked by known agonists (glycine) and antagonists (APV), and detect changes in presynaptic release (following application of phorbol esters). We measure responses to ketamine—currently being investigated for treatment of depression (Berman et al., 2000; Raza et al., 2025), PTSD (Krystal et al., 2017; Valizadeh et al., 2025), acute pain (Gurnani et al., 1996; Trevino et al., 2025), and chronic pain (Carr et al., 2004; Tankha et al., 2025)—and memantine, a preferential antagonist of extrasynaptic NMDARs developed to mitigate the symptoms of advanced Alzheimer’s disease and dementia (Ditzler, 1991) that is now in clinical trials for treatment of Parkinson’s Disease (Jeong & Lee, 2025). We also report effects of and differences between encephalitis patient-derived NMDAR auto-antibodies. By quantifying the function of all active synapses automatically, this pipeline reliably captures a variety of possible effects of compounds on synaptic function in a high-throughput manner.

## Methods

### Cell Culture Preparation

Primary corticohippocampal cultures were prepared under sterile conditions from postnatal day 0 to 2 (P0–P2) wild-type Wistar rat pups using protocols adapted from Turko et al. (2019). Isolated tissue was rapidly dissected and transferred into ice-cold (4°C) cell culture buffer (116 mM NaCl, 5.4 mM KCl, 26 mM NaHCO_3_, 1.3 mM NaH_2_PO_4_, 1 mM MgSO_4_•7H_2_O, 1 mM CaCl_2_•2H_2_O, 0.5 mM EDTA•2Na•2H_2_O, and 25 mM D-glucose, pH = 7.4). Tissue was diced and then incubated at 37°C in 5 mL cell culture buffer containing 1.5 mg/mL Papain (Sigma) for 25 minutes. The digested tissue was then triturated with a fine-tip Pasteur pipette in 3 separate 15 mL Falcon tubes, each containing 4 mL cell culture buffer supplemented with 10 mg/mL bovine serum albumin (BSA; Sigma). The triturated tissue was then combined into a single 12 mL suspension and centrifuged at 3000 RPM for 3 min. The supernatant was discarded, and the pellet resuspended in pre-warmed (37°C) complete Neurobasal-A medium (complete-NBA) without phenol red (Gibco) supplemented with B27 (1x), GlutaMAX (1x), and Penicillin-Streptomycin (100 U/mL; all from Gibco). Cells were then passed through a 30 µm cell strainer (Partec CellTrics; Sysmex). To determine the number of viable cells in suspension, cells were stained with trypan blue and counted on a hemocytometer. Cells were plated at 50,000 cells per well, in 24-well glass-bottom plates with high-performance #1.5H cover glass (0.170±0.005 mm, In Vitro Scientific), in 500 µL of complete-NBA medium without phenol red.

### Viral Transduction

Diluted cell suspensions (1.25 million cells) were treated individually with 1 μL of AAV2/1 human synapsin promoter-driven GCaMP6f (2.0 × 10^13^ v.g./mL, pAAV.Syn.GCaMP6f.WPRE.SV40, Addgene; Chen et al., 2013) and vortexed. Cells were seeded at a density of 50,000 cells in 500 µL complete-NBA medium without phenol red/well (with ∼0.05 µL of virus per well) on poly-L-lysine-coated 24-well glass-bottom plates. Culture plates were maintained inside sealed containers in humidified incubators with 5% CO_2_ at 37°C until experiments were performed.

### Synaptic Calcium Imaging

For synaptic calcium imaging, DIV19-22 rat corticohippocampal cultures were imaged without magnesium (0 mM Mg^2+^) and in the presence of 1 µM tetrodotoxin (TTX) in ACSF (140 mM NaCl, 2.5 mM KCl, 10 mM Glucose, 10 mM HEPES, and 2 mM CaCl_2,_ pH adjusted to 7.4 with NaOH). For overnight treatments, pharmacological agents or antibodies were added to cultures the day before imaging and remained in cultures until media were exchanged for 1 mL of 0 mM Mg^2+^ ACSF with TTX immediately before synapse calcium imaging the next day. For acute treatments, wells were washed once with 0.5 mL of 0 mM Mg^2+^ ACSF with TTX and imaged in 0.5 mL of the same media. Acute treatments were then bath applied in an additional 0.5 mL of 0 Mg^2+^ solution with TTX containing 2× concentrated compound, to promote instantaneous mixing to the desired final concentration.

Cultures were imaged using a Nikon Ti-2 Eclipse widefield inverted fluorescence microscope with an incubation chamber at 32°C and a 40× water-immersion objective (NA 1.25), and 35 ms triggered LED exposure using 20-25% LED power. Calcium transients were recorded at 20 frames per second (fps), i.e., 50 ms interval between images. Regions imaged were manually selected based on neuronal viability, morphological integrity, and visible synaptic calcium transients; these criteria were applied consistently across all conditions. Regions with 1–4 neuronal somata were chosen. A maximum of 9 regions could be automatically imaged in sequence at defined coordinates before the water for immersion had to be replenished. For overnight application of compounds, regions of interest in control or treated conditions were imaged for 3 minutes each. For acute bath application of compounds, regions of interest were imaged for 3 minutes at baseline, pharmacological agents were applied, and the same regions were imaged again for 3 minutes. All compounds tested are summarized in Table 1.

**Table 1:**
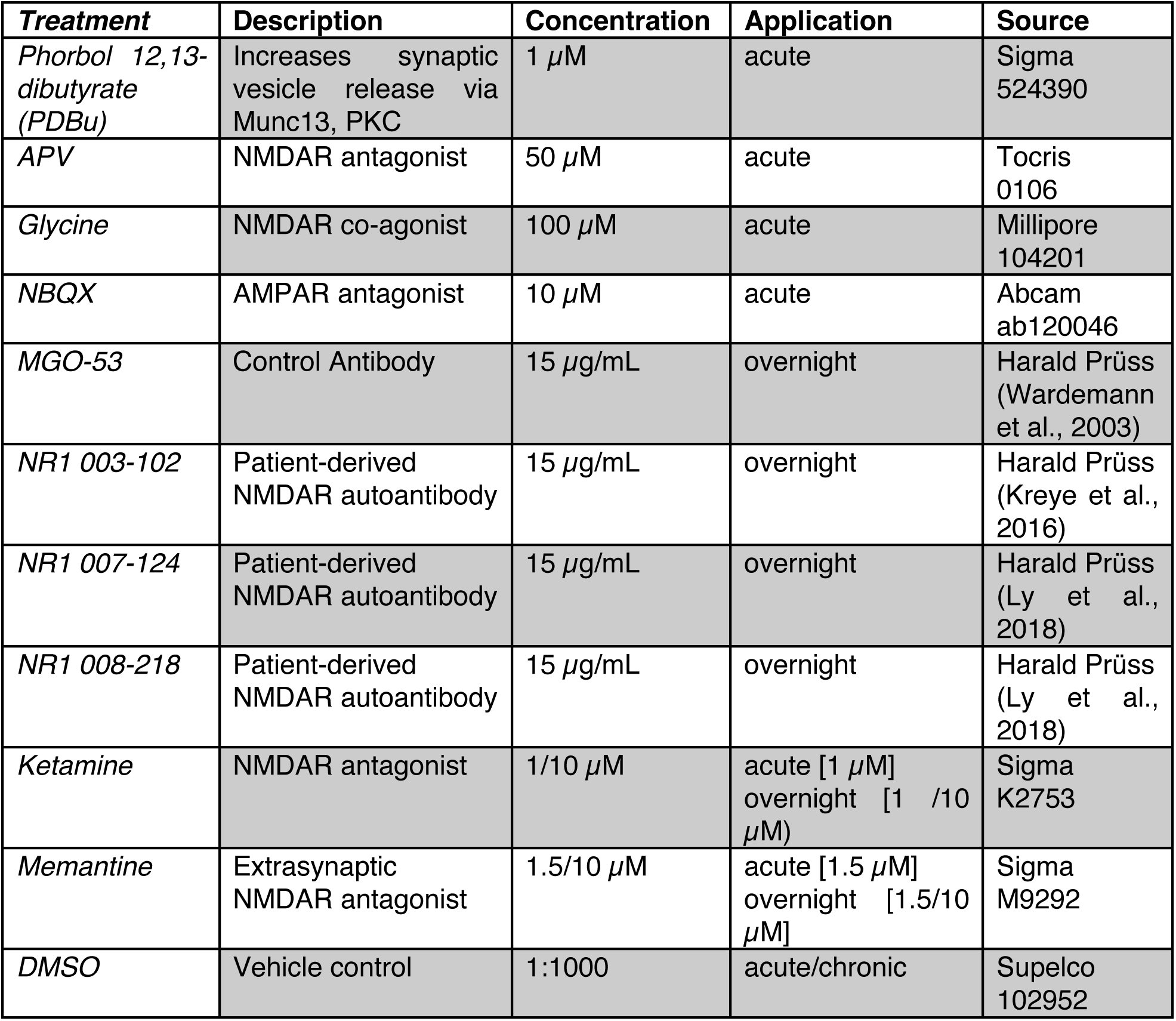
Treatments tested by synapse calcium imaging.

### ROI Detection

Synaptic calcium transients were automatically detected using the calcium imaging analysis tool, Suite2p (Pachitariu et al., 2016). The input parameters for Suite2p were optimized for synapse detection by changing key parameters summarized in Table 2. ROIs detected by Suite2p were automatically filtered to remove noise by excluding those with a fluorescence skew—defined by Suite2p as the deviation of the ROI fluorescence from its neuropil—greater than 1. Only ROIs with a compactness less than or equal to 1.4 (where 1 is a perfect circle) were considered as synaptic calcium transients. More oblong dendritic events were detected but largely excluded from analysis since they were not limited to spines. Both skew and compactness values for each ROI were automatically generated by Suite2p and stored in ‘stat.npy’ files.

**Table 2:** Suite2p ROI detection parameter changes for synaptic calcium imaging.

| **Parameter Category** | **Parameter** | **User input** | **Description** |
| --- | --- | --- | --- |
| *Functional Detect* | sparse_mode | 1.0 | Detect ROIs by searching for changes in fluorescence |
|  | threshold_scaling | 1.5 | Threshold multiplier for ROI inclusion |
|  | connected | 1 | Whether all pixels should be connected in the ROI |
|  | max_overlap | 0.9 | Maximum amount of overlap allowed for ROIs in percent |
| | spatial_scale | 0 | Allows Suite2p to estimate the size of ROIs (for our data, spatial_scale was consistently estimated to be 6 pixels or $\sim 2 \mu\text{m}$ (spatial_scale = 1)) |
| *Extraction/Neuropil* | min_neuropil_pixels | 50 | Minimum number of pixels needed for neuropil fluorescence calculations |
| *Classify/Deconv* | spike_detect | 0 | Turn off deconvolution |
| *Registration* | do_registration | 0 | Stop image registration and motion correction.<br>NOTE: If concatenating multiple videos of the same region, this must be enabled |
| *Output settings* | preclassify | 0 | Removes ROI exclusion by Suite2p classifiers; preserves all detected ROIs |

### Signal Extraction and Normalization

A Python (Python Software Foundation, 2020) pipeline was created to automate preprocessing, Suite2p ROI detection, and analysis of fluorescence of each ROI. After potential ROIs are identified by Suite2p, the pipeline first converted the raw ROI fluorescence and neuropil fluorescence into a normalized change in fluorescence compared to baseline (ΔF/F_0_). To do this, 70% of the neuropil fluorescence was subtracted from the raw fluorescence of each ROI to correct for any potential neuropil contamination. To correct for bleaching and small fluorescence fluctuations, baseline correction was performed using an automated iteratively reweighted least squares regression (Zhang et al., 2010). This method was chosen over other baseline correction methods since it does not require user input parameters to accurately correct baseline values compared to peaks. The airPLS algorithm, used in spectral analyses such as NMR, iteratively measures the difference between its own baseline estimate and the original trace baseline. During the algorithm’s iterations, each point in the trace is reweighed based on the difference between these two values—peaks are given progressively lower weights, while baseline sections are given higher weights—resulting in a flat and corrected baseline.

### Baseline and Calcium Transient Detection

Baseline fluorescence for normalization was determined using median absolute deviation (MAD; Leys et al., 2013). To calculate MAD, the absolute deviation of each data point in the fluorescence trace from the trace median is calculated as:

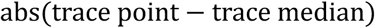

The median of these absolute deviations is then calculated to obtain an initial MAD value. Since the traces are primarily composed of noise, with a few distinct peaks occurring per ROI in each recording, the distribution of fluorescence values for any given calcium trace is right-skewed; however, the noise alone can be considered normally distributed.

The standard deviation is estimated by dividing the trace MAD by the constant 0.6745, specific to normally distributed data. This constant represents the relationship between MAD and the standard deviation (SD) for the middle 50% of a normal distribution, which corresponds to 0.6745 standard deviations. Therefore, the standard deviation (SD) can be calculated as:

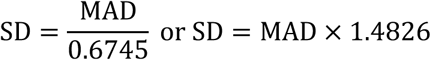

This adjustment allows the standard deviation of a trace to be estimated from the MAD value. Data points with deviations less than 2 standard deviations above the MAD were considered to be noise. These data points without activity were used as a baseline reference (Mitani & Komiyama, 2018).

A refined MAD was then recalculated from the extracted baseline, without confounding peaks that were present in the original MAD calculation, to determine the baseline fluorescence (F_0_) and threshold for detecting peaks. A threshold of 4.5 standard deviations above F_0_ was set empirically based on manual examination of raw data, and used to detect peaks in each baseline-corrected, normalized trace using the ‘find peaks’ function from the SciPy Python package (Virtanen et al., 2020). Constraints on peak detection were set: peaks could only occur greater than or equal to 100 ms (5 frames) apart, peak prominence, or deviation of a peak from local minima, was set to F_0_ + 2 × SD, and the peak half-width was set to be at least 3 frames. The ‘find peaks’ function with these constraints could accurately determine time stamps for when peaks occurred, allowing frequency and inter-event intervals to be calculated. The amplitude of each peak was calculated in relation to F_0_. ROIs with less than 2 detected transients were excluded from frequency analysis. All ROIs with at least one event were included in amplitude and synapse number analyses.

### Normalizing Active Synapses to Neurite Coverage

Cultures treated overnight with compounds and antibodies were compared to untreated control wells. To control for possible differences in neurite coverage, average projection images were first generated automatically using an ImageJ macro. Neurite coverage was measured automatically using CellProfiler v4.2.5 (Stirling et al., 2021). First, soma signals were masked and removed using the ‘Threshold’ and ‘MaskImage’ modules. The signal from neurites was enhanced using the ‘EnhanceOrSuppressFeatures’ and then thresholded and skeletonized using the ‘MorphologicalSkeleton’ module. The length of neurite coverage was calculated using ‘MeasureImageAreaOccupied’ and exported. The thresholded, skeletonized neurites were used to normalize synapse counts. Normalized synapses are reported per 10 µm of neurites.

### Statistical Analysis

In 0 Mg^2+^ and TTX conditions, individual fields of view contained between 1,700 and 3,300 synapses; datasets contained between 150,000 and 340,000 individual synapses. To quantify the effect of pharmacological treatments on synaptic calcium transients, we employed generalized linear mixed-effect models (GLMMs) using R (R Core Team, 2025) and the glmmTMB package (Brooks et al., 2017). This was necessary due to the hierarchical nature of synaptic data. Imaged regions were highly heterogeneous and introduced variability; these were designated as random effects. Other potential random effects, such as culture preparation and well, were not included due to insufficient replicates for accurate modeling (n < 10). Datasets were always compared to relevant control conditions. Synaptic frequency values were analyzed using a generalized linear mixed-effects model fit with glmmTMB in R, where the treatment condition was considered the fixed effect and region as a random effect. Error was assumed to follow a Gamma distribution with a log link:

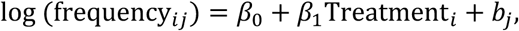

whereas average amplitude was also described using a GLMM, but by assuming a log-normal error distribution for average amplitude values:

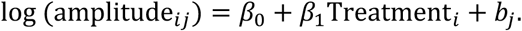

where frequency_PQ_and amplitude_GH_ represent the expected frequency and average amplitude values, respectively, for synapse i in file j, β_J_ is the intercept, β_L_ represents the fixed effect of treatment condition, and b_Q_ is a random intercept accounting for the variability that exists between different regions imaged. Although mathematically similar, the two functions are distinct in assumptions of variance scaling: Gamma distributions assume that variance scales quadratically with the mean, whereas log-normal error distributions assume that the variance scales multiplicatively.

The accuracy of each model for each dataset was confirmed visually using R and DHARMa diagnostics (Hartig, 2024). Quantile-quantile plots were routinely examined to determine if the model accurately predicted simulated data values. Deviations of observed values from expected values, particularly for central values from 0.25 to 0.75 in quantile-quantile plots, indicated potential model misfit. Mixed-effects model results were compared to stratified, clustered bootstrap resampling with replacement, which was repeated 10,000 times. Bootstrap resampling consistently produced similar results to GLMM estimates and effect sizes.

Both mixed-effects models and bootstrap resampling results are visualized using forest plots, which show, on a log scale, the deviation of each treatment group from a reference group. Effect sizes are reported as a fold change relative to a reference group, with 1 being equivalent to the value of controls. Plots show the mean response for frequency or amplitude per group with 95% confidence intervals (CI). Results were considered statistically significant if the confidence interval did not cross one. In this way, true effects could be estimated in the event of model misfit. Wald P-values were generated for each model; however, they were not reported because they are influenced by model assumptions. Analysis and plot generations for mixed-effects models and bootstrap resampling were performed in R. Intraclass correlation coefficients (ICC) indicate the proportion of variance attributable to grouping synapses by imaged region. For each model, marginal and conditional R² values are reported, representing the variance explained by fixed effects and by both fixed and random effects, respectively, together with the ICC.

To compare the number of active synapses before and after acute bath applications, a normality check was performed. For conditions with a singular control and acute treatment condition, if both groups of active synapses passed normality checks, a paired t-test was performed; if not, a Wilcoxon test was performed. Acute and chronic treatments with three or more groups were also checked for normality. Normally distributed synapse counts were analyzed with Friedman tests, while experiments that failed normality checks were analyzed with Kruskal-Wallis tests. Potential outliers were identified in synapse count data using the ROUT method (Q = 5%). Statistical analyses and plot generation for active synapses were done using GraphPad Prism 10 (version 10.5.0). Individual synaptic traces were visualized using matplotlib (Hunter, 2007), and raster plots were generated using pynapple (Viejo et al., 2023).

### Code Availability

The code for synapse analysis in Python, neurite length measurement in CellProfiler, and R mixed-effect models is available here.

## Results

To detect synaptic calcium transients, we transduced corticohippocampal neurons with synapsin promoter-driven GCaMP6f at DIV 0 and imaged between DIV 19 and 22 in 0 magnesium (Mg^2+^), to unblock NMDARs, and TTX to block action potentials. To automatically detect ROIs, we adapted Suite2p (Pachitariu et al., 2016) to detect spontaneous synaptic calcium transients en masse. Synaptic calcium transients were easily identifiable by their localized, large changes in fluorescence relative to baseline during live imaging (<u>Movie S1</u>) and in maximum minus minimum projection images of live imaging over three minutes (Fig. 1A). These spontaneous synaptic calcium transients can be automatically and reliably detected using Suite2p (Fig. 1B) and can be further filtered to remove noise and longer dendritic events (Fig. 1C). Dendritic events spread out from a singular point and last slightly longer than synaptic events, on average, whereas synaptic events most often appear limited to dendritic spines. Example time courses for a single synaptic (Fig. 1D) and dendritic (Fig. 1E) event are shown. Only the punctate, synaptic calcium transients were analyzed, except for a single experiment in which oblong, dendritic events were examined (Fig. 2).

**Figure 1.**
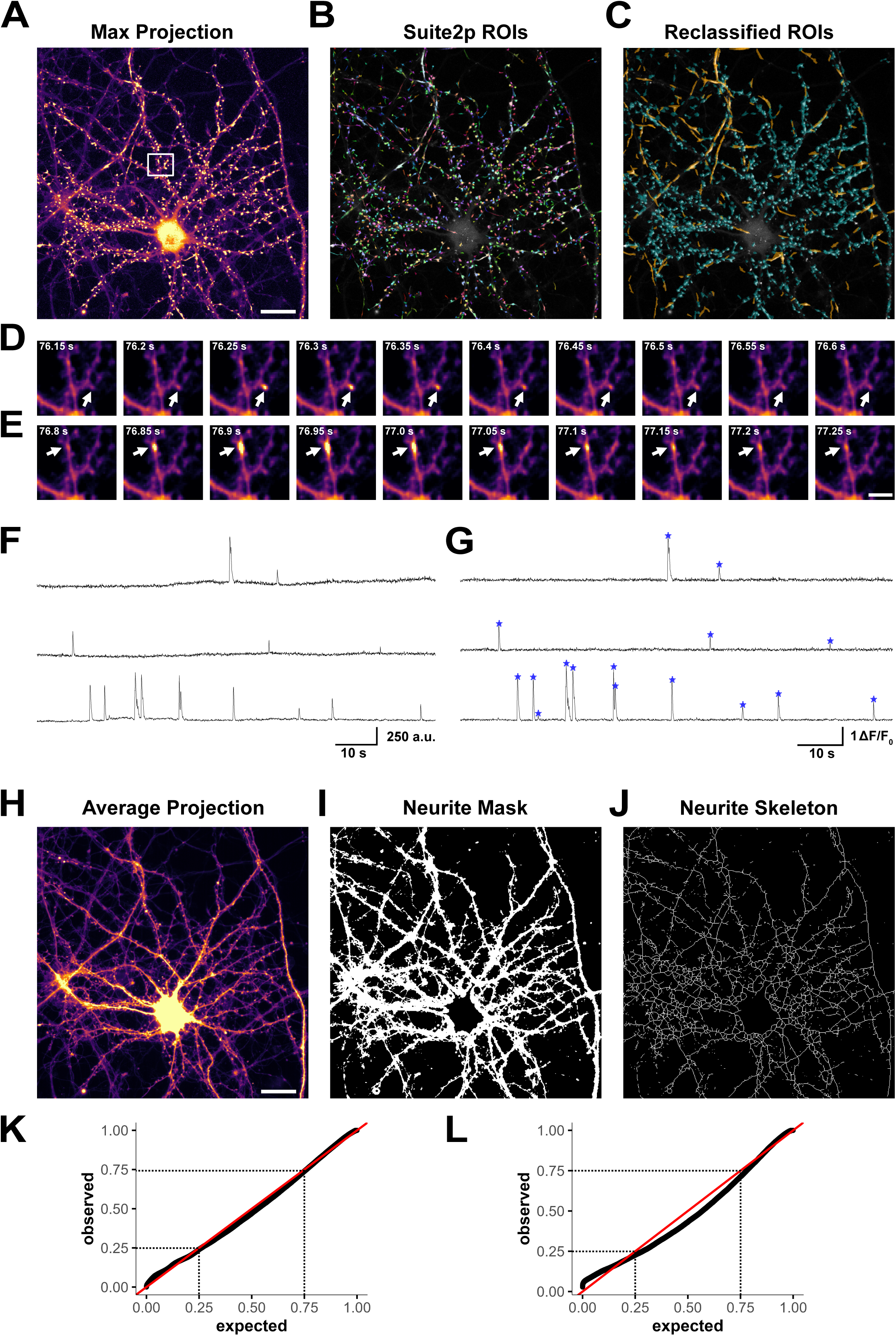
Synaptic calcium imaging analysis can be automated using Suite2p and Python. **A)** Image of maximum fluorescence minus minimum fluorescence projection of a 3-minute calcium imaging recording (scalebar = 25 µm). The region within the white box is expanded in D/E. **B)** Image of maximum fluorescence generated by Suite2p overlayed with Suite2p-detected ROIs with random colors. **C)** Reclassified synaptic (cyan) and dendritic (orange) spontaneous calcium transient ROIs. **D)** Example time course of a representative synaptic calcium transient. **E)** Example time course of a representative dendritic event (scalebar = 5 µm). **F)** Three example raw fluorescence traces from three Suite2p-detected ROIs. **G)** The same three example ROI traces shown in panel F, after airPLS baseline correction and normalization to ΔF/F_0_. Each detected peak is indicated by a blue asterisk. **H)** Average fluorescence projection of a 3-minute calcium imaging recording (scalebar = 25 µm). **I)** Thresholded neurites and masked somatic signal generated by CellProfiler. **J)** Skeletonized neurites generated by CellProfiler were used for normalizing synapse count per length neurite in chronically treated conditions. **K)** Quantile-quantile plot of expected (black) and generated data points (red) for a well-fitting frequency mixed-effects model. **L)** Quantile-quantile plot of expected (black) and generated data points (red) for a poorly-fitting frequency mixed-effects model.

**Figure 2:**
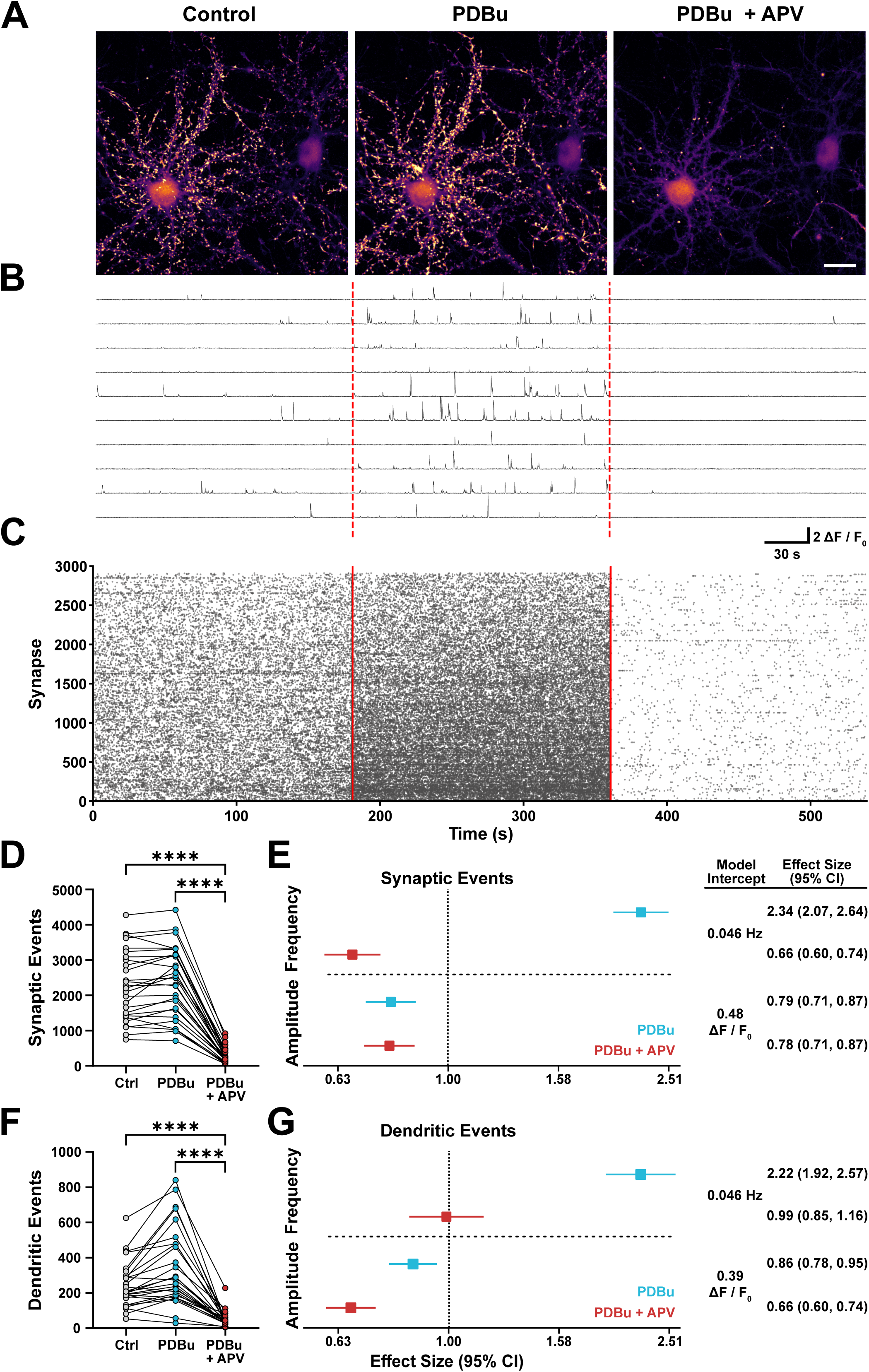
Phorbol esters increase, and APV decreases spontaneous synaptic calcium transient frequency. **A)** Images of maximum fluorescence minus minimum fluorescence projections of 3-minute calcium imaging recordings in control conditions (left), and in the presence of either 1 µM phorbol 12,13-dibutyrate (PDBu; middle) or 1 µM PDBu with 50 µM APV (right; scale bar = 25 µm). **B)** Example concatenated ΔF/F_0_ traces from 10 selected synapses; red dashed lines indicate when each treatment began. **C)** Raster plots showing activity across all synapses in baseline, PDBu, and PDBu plus APV treatment conditions; red solid lines indicate when each treatment began. **D)** Comparisons of the number of spontaneously active synapses in each condition using a Friedman test (n = 57,872 control, 63,777 PDBu-treated, and 5,935 PDBu/APV-treated active synapses). **E)** Forest plot results for synaptic event frequency (top) and amplitude (bottom) obtained from generalized linear mixed-effects models. PDBu (blue) and APV (red) are shown in reference to controls (vertical dashed line). Model intercepts indicate model-predicted averages for frequency or amplitude in control conditions. Effect sizes are reported as mean ± 95% confidence intervals (CI); raw values are summarized on the right. **F)** Comparisons of the number of spontaneously active dendritic events in each condition using a Friedman test (n = 5,881 control, 9,096 PDBu-treated, and 833 PDBu/APV dendritic events). **G)** Forest plot results for synaptic event frequency (top) and amplitude (bottom) obtained from generalized linear mixed-effects models. PDBu (blue) and APV (red) are shown in reference to controls (vertical dashed line). Model intercepts indicate model-predicted averages for frequency or amplitude in control conditions. Effect sizes are reported as mean ± 95% CI; raw values are summarized on the right. **Model Metrics**: Synapse frequency GLMM: ICC: 0.11, Marginal R^2^: 0.30, Conditional R^2^: 0.38. Synapse amplitude GLMM: ICC: 0.15, Marginal R^2^: 0.06, Conditional R^2^: 0.20. Dendritic event frequency GLMM: ICC: 0.12, Marginal R^2^: 0.22, Conditional R^2^: 0.32. Dendritic event amplitude GLMM: ICC: 0.13, Marginal R^2^: 0.05, Conditional R^2^: 0.17 (3 culture preparations; 9 wells imaged; 27 regions; DIV 19) * p<0.05, ** p<0.01, *** p<0.001, **** p<0.0001

The raw fluorescence extracted from Suite2p for each ROI (Fig. 1F) was then baseline corrected using an automated iterative least squares regression (airPLS; Zhang et al., 2010) and normalized (ΔF/F_0_) (Fig. 1G). Calcium transients were detected using the scipy ‘find_peaks’ function and an empirically determined threshold of 4.5 standard deviations above F_0_ for peak detection (Fig. 1G). Functional synapse number before and after treatment was quantified within the same imaged region in experiments with acute treatments. In experiments with overnight treatments, to account for potential differences in neuron/neurite coverage between randomly selected regions, we normalized functional synapse counts to neurite coverage, determined using GCaMP6f average projection images (Fig. 1H) and CellProfiler pipelines. We removed somatic signal and created a binary mask of the neurites (Fig. 1I). This mask was skeletonized (Fig. 1J), and the number of synapses was normalized to the length of neurite coverage. Frequency and amplitude data were analyzed using mixed-effects models. Model accuracy was assessed using quantile-quantile plots that showed well-fitting (Fig. 1K) or poorly-fitting (Fig. 1L) mixed-effects models. All amplitude models were well-fitting whereas some frequency models were poorly-fitting. Results from poorly-fitting models were verified using bootstrap resampling.

### Synaptic calcium transients increase in response to phorbol esters and are blocked by APV

To test if changes in functional synapse number and presynaptic release probability could be assayed using our automated analysis pipeline, we bath-applied the phorbol ester phorbol 12,13-dibutyrate (PDBu, 1 µM). PDBu interferes with Munc13 and protein kinase C (PKC) in the presynaptic terminal, which leads to an increase in presynaptic vesicle release probability (Lou et al., 2008). We then applied APV (50 µM) in sequence following phorbol ester application, to test whether synaptic transients are NMDA-dependent. APV is a preferential, potent, and well-established competitive antagonist of NMDARs (Collingridge et al., 1983).

Compared to control conditions, application of PDBu increased, and subsequent APV treatment decreased, synaptic fluorescence signal in maximum minus minimum projection images of all active synapses recorded during each treatment phase (Fig. 2A; <u>Movie S2</u>). Individual fluorescence traces showed changes in frequency across the three conditions at individual synapses (Fig. 2B) and raster plots showed that these patterns persisted across all detected synapses (Fig. 2C). PDBu did not change the number of active synapses compared to controls (∼2,398 synapses; p = 0.40). APV addition decreased the number of active synapses approximately seven-fold (from ∼2,268 synapses to ∼319 synapses; p < 0.0001) compared to both control and PDBu-treated conditions (Fig. 2D).

In control conditions, the average frequency was estimated to be 0.046 Hz. PDBu bath application acutely increased frequency to 0.11 Hz (2.34, 95% CI 2.07–2.64). APV decreased the frequency to 66% of control levels (0.031 Hz, 95% CI 0.60–0.74; Fig. 2E, top). The average amplitude of synaptic calcium transients at baseline was 0.48 ΔF/F_0_ in control conditions. Average amplitude decreased in the presence of PDBu to 0.38 ΔF/F_0_, 79% of control (0.79, 95% CI 0.71–0.87). Subsequent addition of APV did not further decrease average amplitude compared to controls (0.78, 95% CI 0.71–0.87; Fig. 2E, bottom). Together, these findings illustrate that synaptic calcium transients are dependent on presynaptic glutamate release and driven by NMDAR activity.

Dendritic calcium transients were also analyzed in control and PDBu and APV-treated conditions. Like synaptic events, dendritic calcium transients dramatically decreased in number in the presence of APV compared to both control and PDBu-treated conditions (p < 0.0001, Fig. 2F). These events also responded to PDBu application with an increase in event frequency at individual synapses: at baseline, average dendritic event frequency for individual event sites was 0.046 Hz, and PDBu application increased frequency to 0.10 Hz (2.22, 95% CI 1.92–2.57). Interestingly, although APV application blocked ∼82% of dendritic events, the remaining active sites had almost identical frequencies to baseline (0.99, 95% CI 0.85–1.16; Fig. 2G, top). The amplitude of dendritic events was decreased in the presence of PDBu (0.86, 95% CI 0.78–0.95); however, dendritic event amplitude was then further decreased with the addition of APV (0.66, 95% CI 0.60–0.74), unlike synaptic events. Since dendritic events responded differently to APV than expected, they were not included in further analysis of NMDAR events, where we focused on synaptic events.

### Glycine increases synaptic calcium transient frequency and number

To further test the NMDAR-dependence of synaptic calcium transients, we bath-applied the NMDAR co-agonist glycine at a concentration of 100 µM to cultures. This concentration is saturating for NMDARs (Cummings & Popescu, 2015). Glycine led to an immediate increase in fluorescence, which can be seen in maximum minus minimum projection images (Fig. 3A). Traces from individual synapses (Fig. 3B) and raster plots across all synapses (Fig. 3C) revealed a dramatic increase in frequency of synaptic calcium transients in the presence of glycine. Glycine significantly increased the number of active synapses from ∼2,503 to ∼2,832 (p < 0.0001; Fig. 3D). Furthermore, the frequency of synaptic calcium transient events increased by 55% from 0.051 Hz to 0.079 Hz (1.55, 95% CI 1.38–1.74; Fig. 3E, top). The average amplitude of synaptic calcium transients simultaneously decreased by 15% from 0.50 to 0.43 ΔF/F_0_ (0.85, 95% CI 0.78–0.93; Fig. 3E, bottom). The increase in number of active synapses and event frequency in response to glycine illustrates that synaptic calcium imaging reports multiple actions of NMDAR agonists, including postsynaptic changes in NMDA-receptor function and increases in functional synapse number.

**Figure 3:**
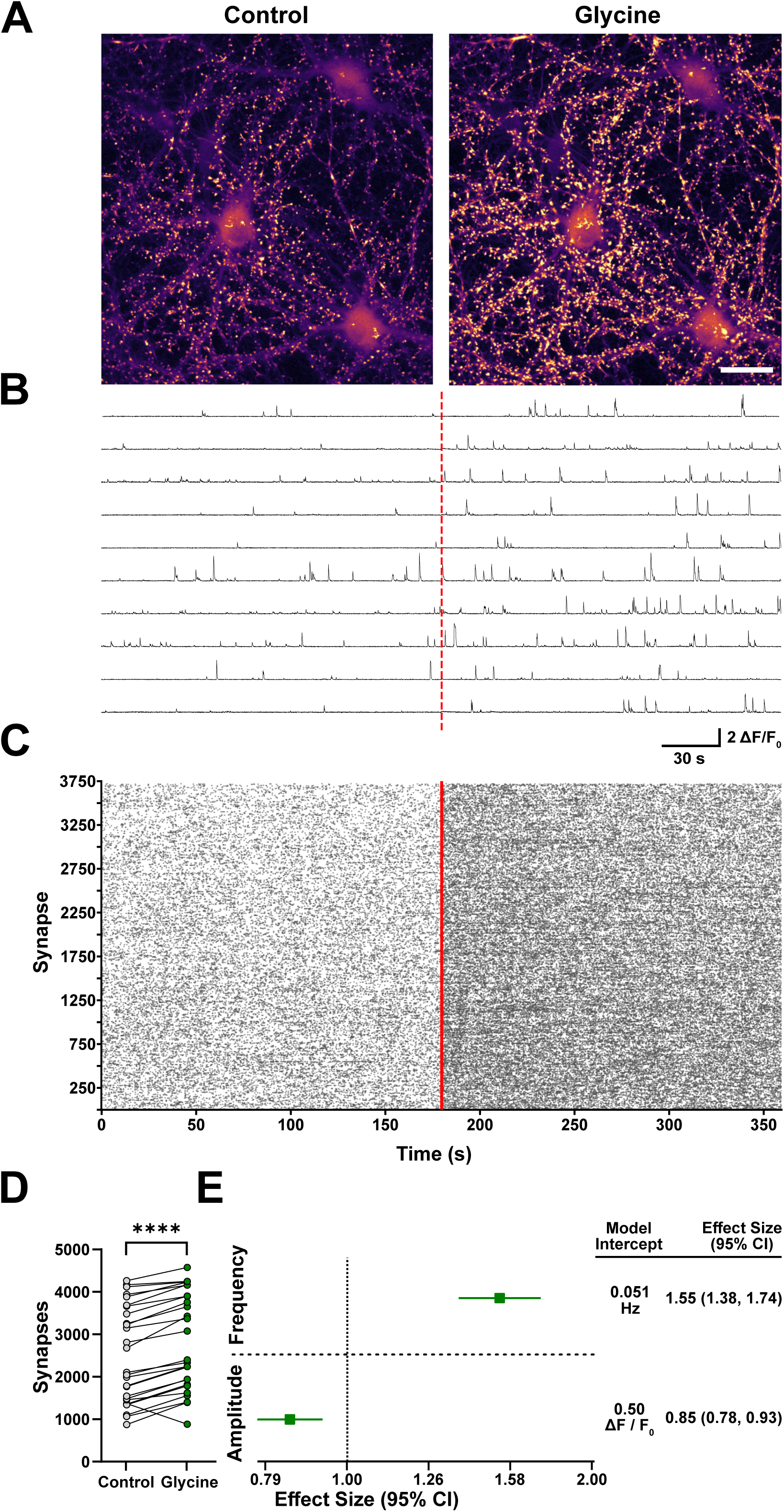
Glycine increases spontaneous synaptic calcium transient frequency and number of active synapses. **A)** Images of maximum fluorescence minus minimum fluorescence projections of 3-minute calcium imaging recordings in control conditions and in the presence of 100 µM glycine (right; scale bar = 25 µm). **B)** Example concatenated ΔF/F_0_ traces from 10 selected synapses; the red dashed line indicates when treatment began. **C)** Raster plots of activity across all synapses at baseline and with glycine treatment; the red solid line indicates when treatment began. **D)** Comparisons of the number of spontaneously active synapses in each condition using a Wilcoxon Signed-Rank Test (n = 64,788 control, 75,509 glycine-treated active synapses). **E)** Forest plot results for synaptic event frequency (top) and amplitude (bottom) obtained from generalized linear mixed-effects models. Glycine (green) is shown in reference to controls (vertical dashed line). Model intercepts indicate model-predicted averages for frequency or amplitude in control conditions. Effect sizes are reported as mean ± 95% CI; raw values are summarized on the right. **Model Metrics**: Frequency GLMM: ICC = 0.11, Marginal R² = 0.11, Conditional R² = 0.21. Amplitude GLMM: ICC = 0.12, Marginal R² = 0.03, Conditional R² = 0.14. (3 culture preparations; 9 wells; 27 regions; DIV 19). * p<0.05, ** p<0.01, *** p<0.001, **** p<0.0001

### AMPA receptors have limited contribution to synaptic calcium transients

To test if AMPA receptors (AMPARs) contribute to synaptic calcium transients, we bath-applied the AMPAR antagonist NBQX at a concentration of 10 µM. A general increase in fluorescence of synaptic events appeared following NBQX treatment (Fig. 4A). An increase is also seen in the amplitude of individual events in single synaptic traces (Fig. 4B), but no difference in event frequency is apparent in raster plots (Fig. 4C). Comparison of synapse number between control and NBQX-treated conditions showed no significant difference, although there was a trend toward an increase in synapse number from ∼2,331 to ∼2,401 within each imaged region before and after treatment (Fig. 4D, p = 0.088).

**Figure 4:**
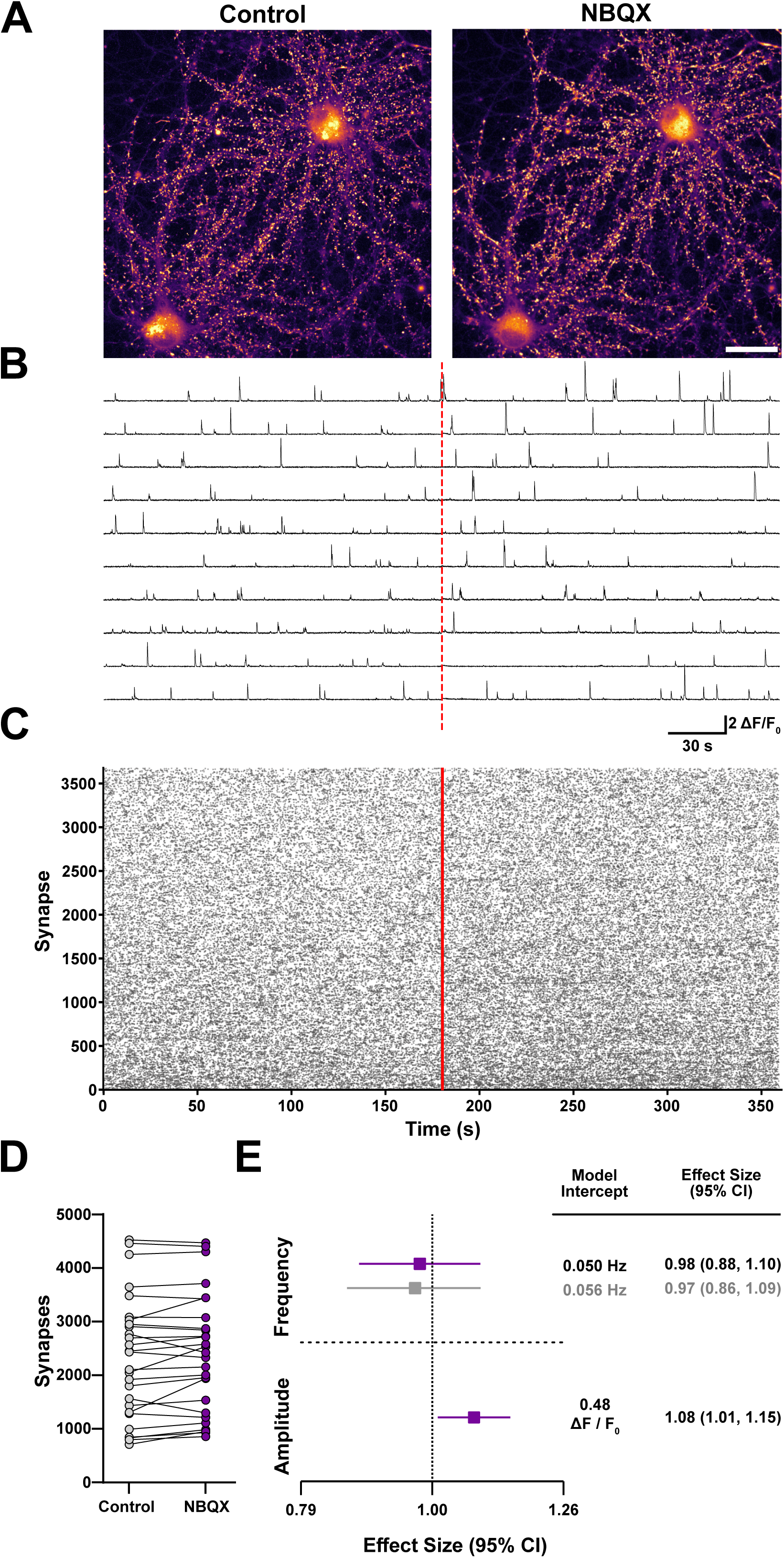
AMPARs show limited contribution to spontaneous synaptic calcium transients. **A)** Images of maximum fluorescence minus minimum fluorescence projections of 3-minute calcium imaging recordings in control conditions and in the presence of 10 µM NBQX (right; scale bar = 25 µm). **B)** Example concatenated ΔF/F_0_ traces from 10 selected synapses; the red dashed line indicates when treatment began. **C)** Raster plots of activity across all synapses at baseline and with glycine treatment. **D)** Comparisons of the number of spontaneously active synapses in each condition using a paired t-test (n = 59,770 control, 61,988 NBQX-treated active synapses). **E)** Forest plot results for synaptic event frequency (top) and amplitude (bottom) obtained from generalized linear mixed-effects models. NBQX (purple) is shown in reference to controls (vertical dashed line). The gray point in the frequency forest plot indicates the estimated mean ± 95% CI from bootstrap resampling (n=10,000). Model intercepts indicate model-predicted averages for frequency or amplitude in control conditions from mixed-effects models (black) or bootstrap resampling (gray). Effect sizes are reported as mean ± 95% CI; raw values are summarized on the right for mixed-effects models (black) and bootstrap resampling (gray). **Model Metrics**: Frequency GLMM: ICC = 0.09, Marginal R² = 0.00, Conditional R² = 0.09. Amplitude GLMM: ICC = 0.06, Marginal R² = 0.01, Conditional R² = 0.06. (3 culture preparations; 9 wells imaged; 27 regions; DIV 19) * p<0.05, ** p<0.01, *** p<0.001, **** p<0.0001

NBQX application had no significant effect on the frequency of synaptic events (0.98, 95% CI 0.88–1.10; Fig. 4E, top). Bootstrap resampling supported GLMM results, verifying that NBQX application did not alter the frequency of synaptic calcium transients (Fig. 4E, top; grey). Inhibition of AMPARs did, however, significantly increase the average amplitude of synaptic calcium transients from 0.48 ΔF/F_0_ to 0.52 ΔF/F_0_ (1.08, 95% CI 1.01–1.15). This increase likely occurs because more glutamate is available to bind NMDARs in the presence of NBQX.

### Acute effects of NMDAR-targeting compounds: ketamine and memantine reduce synaptic activity

After demonstrating that synaptic calcium imaging can report increases and decreases in numbers of functional synapses and changes in presynaptic release, in addition to postsynaptic effects, we next sought to test the potential of this pipeline for screening the NMDAR-targeting compounds ketamine and memantine. These compounds were tested for acute and overnight effects. The IC_50_ for ketamine and memantine were measured in rat hippocampal neuronal culture using electrophysiology to be 0.43 ± 0.10 µM and 1.04 ± 0.26 µM, respectively (Parsons et al., 1996). We initially decided to test concentrations slightly higher than this: 1 µM for ketamine and 1.5 µM for memantine.

Ketamine or memantine was bath applied to cultures in 0 mM Mg^2+^ and TTX acutely. In maximum minus minimum fluorescence projection images, a marked decrease in the number of active synapses was observed for both compounds (Fig. 5A). Fluorescence traces of individual synapses also showed a decrease in the frequency and amplitude of synaptic calcium transients after treatment compared to baseline (Fig. 5B). In raster plots we also observed a decrease in the density of synaptic calcium transients in treated conditions compared to control conditions across all synapses (Fig. 5C). Both ketamine and memantine decreased the total number of active synapses. Ketamine decreased the number of active synapses from ∼2,179 at baseline to ∼1,769 after treatment (p < 0.0001; Fig. 5D, left) while memantine decreased the number of active synapses from ∼2,376 to ∼1,427 (p < 0.0001; Fig. 5D, right). Acute application of DMSO vehicle did not significantly change the number of active synapses (baseline: ∼1,800, treated: ∼1,868, p = 0.40; data not shown).

**Figure 5:**
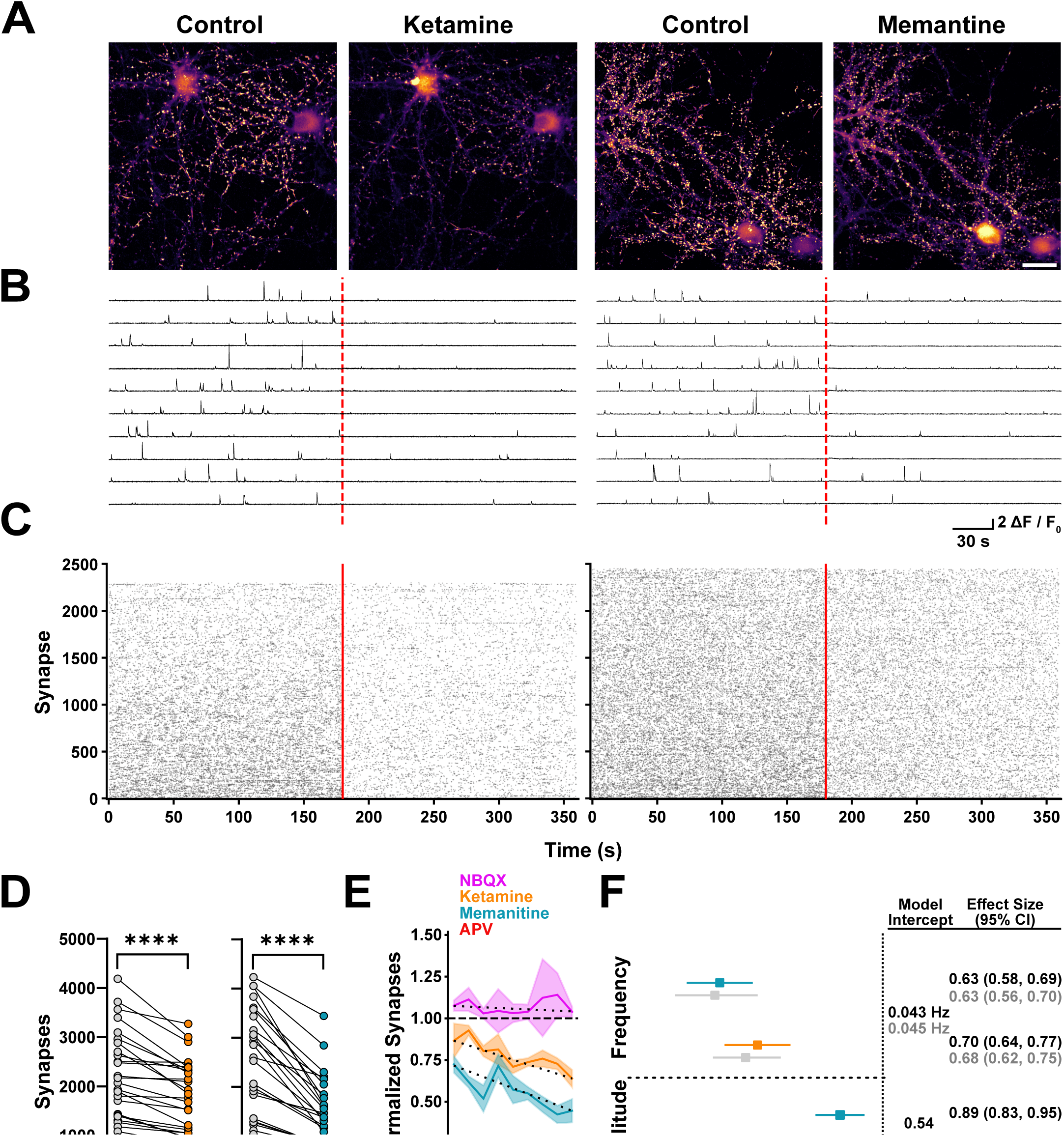
Ketamine and memantine decrease spontaneous synaptic calcium transient frequency and amplitude. **A)** Images of maximum fluorescence minus minimum fluorescence projections of 3-minute calcium imaging recordings in control conditions and in the presence of either 1 µM ketamine (left) or 1.5 µM memantine (right; scale bar = 25 µm). **B)** Example concatenated ΔF/F_0_ traces from 10 randomly selected synapses; the red dashed lines indicate when each treatment began. **C)** Raster plots of activity across all synapses at baseline and with either ketamine (left) or memantine (right) treatment; the red solid lines indicate when each treatment began. **D)** Comparisons of the total number of spontaneously active synapses in control (n = 55,397) and ketamine conditions (n = 42,691) using a paired t-test (left) and active synapses in control (n = 61,143) and memantine (n = 32,548) conditions using a Wilcoxon Signed-Rank Test (right). **E)** Line plots showing the proportion of active synapses after treatment, normalized to active synapses at baseline in the same region. **F)** Forest plot results for synaptic event frequency (top) and amplitude (bottom) obtained from generalized linear mixed-effects models. Ketamine (orange) and memantine (cyan) are shown in reference to DMSO vehicle controls (vertical dashed line). Bootstrap results for each treatment group frequency (gray) are shown directly below the mixed-effects model results. Model intercepts indicate model-predicted averages for frequency or amplitude in control conditions from mixed-effects models (black) or bootstrap resampling (gray). Effect sizes are reported as mean ± 95% CI; raw values are summarized on the right for mixed-effects models (black) and bootstrap resampling (gray). **Model Metrics**: Frequency GLMM: ICC = 0.08, Marginal R² = 0.10, Conditional R² = 0.17. Amplitude GLMM: ICC = 0.06, Marginal R² = 0.01, Conditional R² = 0.07. (3 culture preparations; 9 wells imaged; 27 regions; DIV 19) * p<0.05, ** p<0.01, *** p<0.001, **** p<0.0001

Imaging multiple regions in sequence enabled us to track ketamine and memantine antagonism of NMDARs over time. Percentages of active synapses after treatment were normalized to the number of active synapses in the same region at baseline. Each image, in sequence, allowed us to track how the proportion of active synapses changed over time after acute bath applications. We observed that ketamine and memantine progressively increase effects on synapses over time, whereas APV and NBQX produce immediate and stable effects over 30 minutes. (Fig. 5E). Following ketamine treatment, 88.8 ± 17.8% of synapses were active in the first 2–3 minutes after application, and 63.9 ± 9.0% of synapses were active in the last imaged region 30 minutes later (orange). In the first 2–3 minutes after memantine application, the proportion of active synapses compared to baseline was 72.4 ± 8.7%, and in the last region imaged, the proportion dropped to 44.7 ± 12.1%. The decrease in active synapses over time is supported by previous evidence that both ketamine and memantine antagonism of NMDARs only occurs after channel opening (H. Chen et al., 1992; MacDonald et al., 1987). Memantine is also known to have faster blocking kinetics than ketamine (Parsons et al., 1995), although ketamine has a higher affinity for NMDARs. APV antagonism has faster effects, with only 13.8 ± 4.7% active synapses remaining immediately after acute application and 9 ± 4.8% active synapses approximately 30 minutes later. Conversely, NBQX addition increased synapse number immediately to 108 ± 5.4%, and this increase was maintained until the last imaging session, with 104 ± 5.2% of baseline synapses after approximately 30 minutes. Rather than imaging a single region for 30 minutes to track inhibition of individual synapses over time, in multiple regions imaged in sequence we can detect a decrease in the number of active synapses over time, after acute treatment of ketamine or memantine, relative to a baseline reference.

Compared to controls, both ketamine and memantine decreased synaptic calcium transient frequency. The frequency in DMSO vehicle-treated control conditions was 0.043 Hz. Ketamine decreased frequency to 0.030 Hz (0.70, 95% CI 0.64–0.77) and memantine decreased frequency to 0.027 Hz (0.63, 95% CI 0.58–0.69) (Fig. 5F, top). Bootstrap resampling supported frequency model results with similar effects observed in each (Fig. 5F, top; grey). The average amplitude of synaptic calcium transients in DMSO controls was 0.54 ΔF/F_0_. Acute ketamine treatment trended toward decreased average amplitude (0.93, 95% CI 0.87–1.00) while memantine decreased average amplitude to 0.48 ΔF/F_0_ (0.89, 95% CI 0.83–0.95; Fig. 5F, bottom). DMSO vehicle did not change the frequency or amplitude of synaptic events acutely (data not shown).

### Chronic ketamine treatment reduces the frequency and amplitude of synaptic calcium transients

Since ketamine is tested clinically for its antidepressant actions, we also tested what synaptic changes could occur with overnight exposures of cultures to ketamine. Ketamine was applied overnight at 1 µM and an order of magnitude more, at 10 µM. Cultures remained in ketamine until they were switched into a 0 Mg^2+^ TTX-containing solution without ketamine for synaptic calcium imaging. Maximum minus minimum projection images (Fig. 6A) and synaptic transient fluorescence traces (Fig. 6B) showed a concentration-dependent decrease in synaptic calcium transient amplitude in synapses treated overnight with ketamine. Raster plots of activity across all detected synapses hint at both a decreased density of events and fewer synapses in regions treated with 10 µM ketamine overnight compared to control and 1 µM ketamine treatment (Fig. 6C). Interestingly, overnight ketamine treatment did not significantly change synapse number per 10 µm of neurites compared to DMSO vehicle controls (1 µM ketamine: p = 0.91, 10 µM ketamine: p = 0.69). However, there was a concentration-dependent effect of overnight ketamine treatment, with 10 µM ketamine leading to less active synapses compared to 1 µM ketamine (p = 0.023, Fig. 6D).

**Figure 6:**
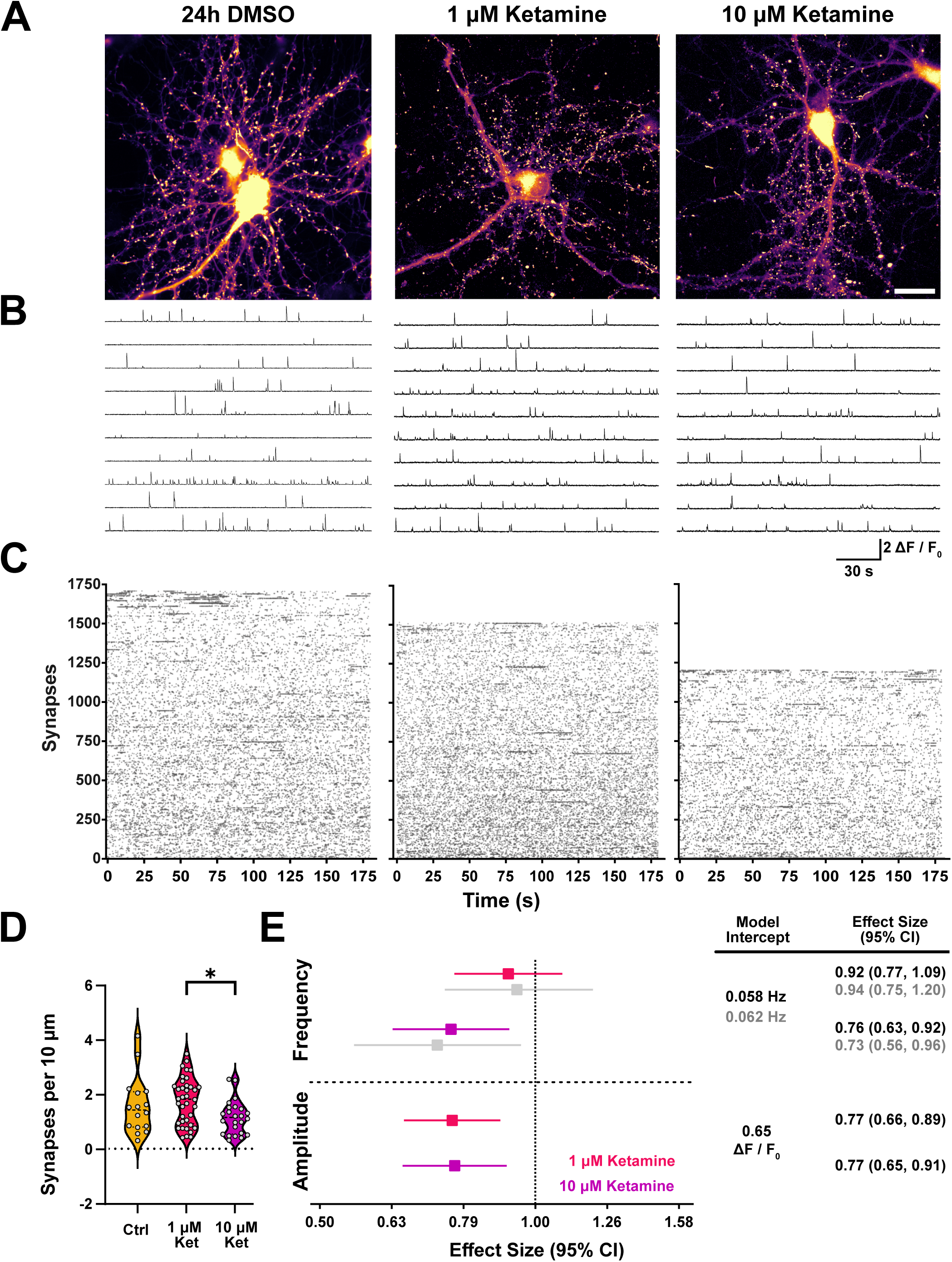
Overnight ketamine treatment reduces the frequency and amplitude of synaptic events. **A)** Images of maximum fluorescence minus minimum fluorescence projections of 3-minute synaptic calcium imaging recordings of regions treated overnight with a DMSO vehicle (left), 1 µM ketamine (middle), or 10 µM ketamine (right; scale bar = 25 µm). **B)** Example ΔF/F_0_ traces from 10 randomly selected synapses from each condition. **C)** Raster plots of activity across all synapses in each treatment condition. **D)** Comparisons of the number of spontaneously active synapses in each condition per 10 µm of GCaMP6f neurite coverage using a Kruskal-Wallis test. **E)** Forest plot results for synaptic event frequency (top) and amplitude (bottom) obtained from generalized linear mixed-effects models. 1 µM ketamine (red) and 10 µM ketamine (purple) are shown in reference to 24h DMSO vehicle controls (vertical dashed line). Bootstrap results for each ketamine treatment for frequency (gray) are shown directly below mixed-effects model results. Model intercepts indicate model-predicted averages for frequency or amplitude in control conditions from mixed-effects models (black) or bootstrap resampling (gray). Effect sizes are reported as mean ± 95% CI, raw values are summarized on the right for mixed-effects models (black) and bootstrap resampling (gray). **Model Metrics**: Frequency GLMM: ICC = 0.12, Marginal R² = 0.03, Conditional R² = 0.14. Amplitude GLMM: ICC = 0.19, Marginal R² = 0.02, Conditional R² = 0.20. (3-5 culture preparations; 15–38 regions; DIV 19 and DIV 22) * p<0.05, ** p<0.01, *** p<0.001, **** p<0.0001

Overnight treatment with DMSO vehicle produced an average frequency of 0.058 Hz. Only 10 µM ketamine significantly decreased calcium transient frequency to 0.044 Hz (0.76, 95% CI 0.63–0.92; Fig. 6E top). Mixed-effects results were compared to bootstrap resampling, which showed similar effects (Fig. 6E top, grey). The amplitude of synaptic calcium transients was also compared between DMSO and the two concentrations of ketamine. Interestingly, both 1 µM and 10 µM ketamine decreased synaptic calcium transient amplitude equally from 0.65 ΔF/F_0_ to 0.50 ΔF/F_0_ (1 µM ketamine: 0.77, 95% CI 0.66–0.89; 10 µM ketamine: 0.77, 95% CI 0.65–0.91; Fig. 6E, bottom).

### Chronic memantine treatment reduces synaptic calcium transient amplitude

Like ketamine, we also tested overnight treatment of memantine at 1.5 µM and 10 µM. Maximum minus minimum projection images showed a decrease in fluorescence of synaptic calcium transients in regions treated with 10 µM memantine compared to controls and 1.5 µM memantine (Fig. 7A). Traces from individual synapses suggested a concentration-dependent decrease in synaptic calcium transient amplitude with increasing concentrations of memantine (Fig. 7B). Raster plots of total activity did not reveal any obvious changes in frequency of events (Fig. 7C). Overnight memantine treatment did not change the number of active synapses (Fig. 7D), or synaptic calcium transient frequency, at either concentration; 1.5 µM memantine trended toward increasing frequency (1.11, 95% CI 0.93–1.33) while 10 µM trended toward decreasing frequency (0.91, 95% CI 0.75–1.10) (Fig. 7E, top). Bootstrap resampling supported model results of no significant difference in frequency, and the trend in difference in effect direction with 1.5 µM and 10 µM memantine (Fig. 7E, top; grey). Treatment with 10 µM memantine decreased the amplitude of calcium transients compared to controls (0.80, 95% CI 0.68–0.95; Fig 8E, bottom). The effect of memantine on synaptic calcium transients is weaker overnight compared to acute applications in 0 Mg^2+^. This is likely because memantine binds NMDARs more weakly in the presence of 1 mM Mg^2+^ (Kotermanski & Johnson, 2009), which was present during overnight treatment in Neurobasal-A medium.

**Figure 7:**
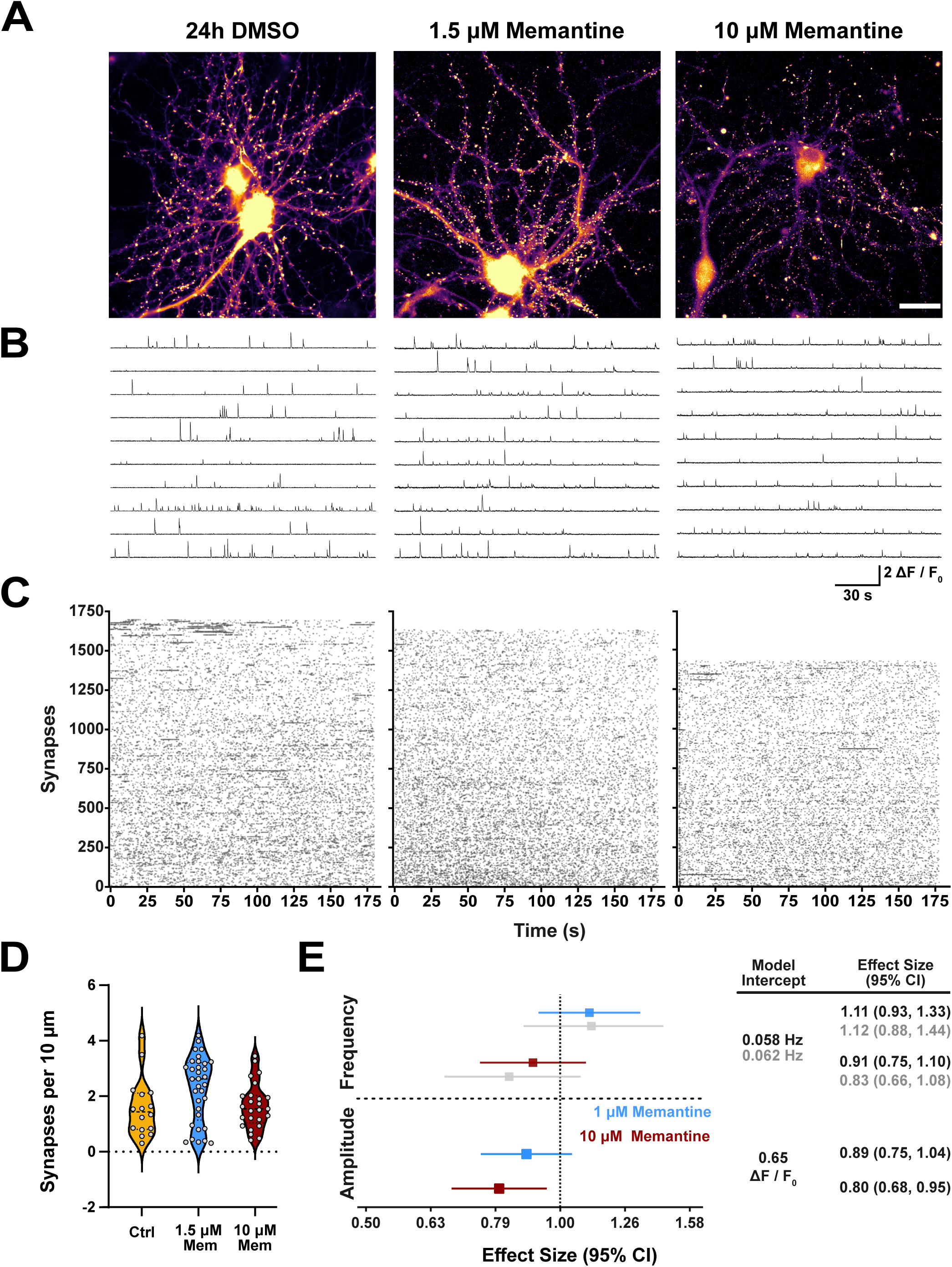
Overnight memantine treatment decreases the amplitude of synaptic events. **A)** Images of maximum fluorescence minus minimum fluorescence projections of 3-minute synaptic calcium imaging recordings of regions treated overnight with a DMSO vehicle (left), 1.5 µM memantine (middle), or 10 µM memantine (right; scale bar = 25 µm). **B)** Example ΔF/F_0_ traces from 10 randomly selected synapses from each condition. **C)** Raster plots of activity across all synapses in each treatment condition. **D)** Comparisons of the number of spontaneously active synapses in each condition per 10 µm of GCaMP6f neurite coverage using a Kruskal-Wallis test. **E)** Forest plot results for synaptic event frequency (top) and amplitude (bottom) obtained from generalized linear mixed-effects models. 1.5 µM memantine (blue) and 10 µM memantine (burgundy) are shown in reference to 24h DMSO vehicle controls (vertical dashed line). Bootstrap results for each memantine treatment for frequency (gray) are shown directly below mixed-effects model results. Model intercepts indicate model-predicted averages for frequency or amplitude in control conditions from mixed-effects models (black) or bootstrap resampling (gray). Effect sizes are reported as mean ± 95% CI, raw values are summarized on the right for mixed-effects models (black) and bootstrap resampling (gray). **Model Metrics**: Frequency GLMM: ICC = 0.12, Marginal R² = 0.03, Conditional R² = 0.14. Amplitude GLMM: ICC = 0.19, Marginal R² = 0.02, Conditional R² = 0.20. (3-5 culture preparations; 15–29 regions; DIV 19 and DIV 22) * p<0.05, ** p<0.01, *** p<0.001, **** p<0.0001

**Figure 8:**
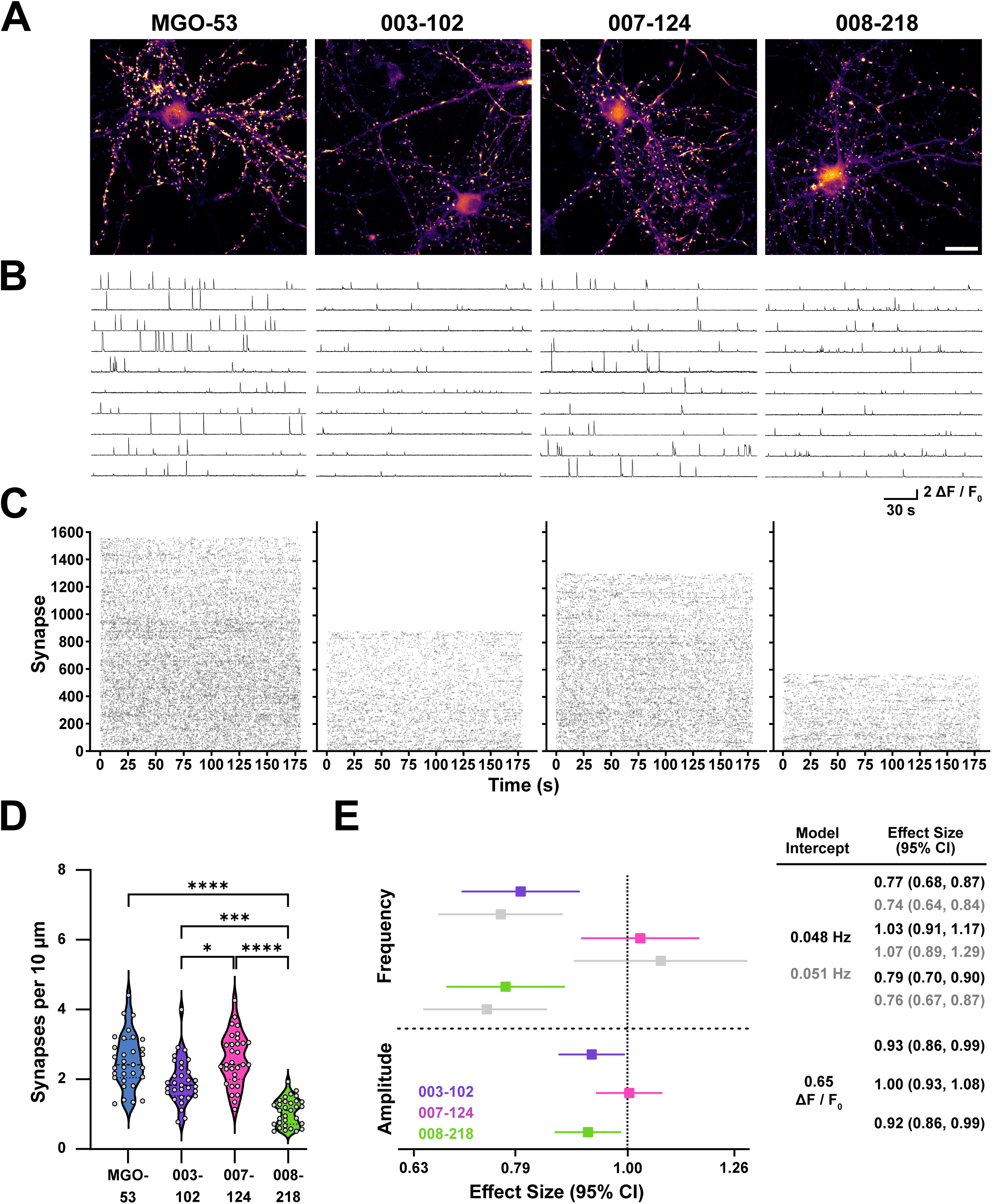
Encephalitis patient-derived NMDAR autoantibodies reduce the number of active synapses and impair synaptic calcium transient frequency and amplitude. **A)** Images of maximum fluorescence minus minimum fluorescence projections of 3-minute synaptic calcium imaging recordings of regions treated overnight with 15 µg/µL of either control antibody (MGO-53, left), or patient-derived NMDAR autoantibodies (003-102, 007-124, and 008-218; right; scale bar = 25 µm). **B)** Example ΔF/F_0_ traces from 10 selected synapses from each condition. **C)** Raster plots of activity across all synapses in each treatment condition. **D)** Comparisons of the number of spontaneously active synapses in each condition per 10 µm of GCaMP6f neurite coverage using the non-parametric Kruskal-Wallis test. **E)** Forest plot results for synaptic event frequency (top) and amplitude (bottom) obtained from generalized linear mixed-effects models. Antibody treatments 003-102 (purple), 007-124 (pink), and 008-218 (green) are shown in reference to MGO-53 controls (vertical dashed line). Bootstrap results for each treatment group frequency are shown in gray directly below the mixed-effects model results. Model intercepts indicate model-predicted averages for frequency or amplitude in control conditions from mixed-effects models (black) or bootstrap resampling (gray). Effect sizes are reported as mean ± 95% CI; raw values are summarized on the right for mixed-effects models (black) and bootstrap resampling (gray). **Model Metrics**: Frequency GLMM: ICC = 0.12, Marginal R² = 0.03, Conditional R² = 0.14. Amplitude GLMM: ICC = 0.06, Marginal R² = 0.005, Conditional R² = 0.06. (4 culture preparations; 15 wells imaged; 31 regions; DIV 20) * p<0.05, ** p<0.01, *** p<0.001, **** p<0.0001

### Synaptic calcium imaging detects differential effects of patient-derived NMDAR autoantibodies

Lastly, we investigated three encephalitis patient-derived anti-NMDAR autoantibodies to test whether our pipeline could detect differences in effects on synapses. The antibodies tested were 003-102, 007-124, and 008-218 (Kreye et al., 2016; Ly et al., 2018) compared to MGO-53 as a negative control (Kreye et al., 2016; Michalski et al., 2024; Wardemann et al., 2003). MGO-53 is a non-reactive patient antibody from mature, naïve B-cells (Wardemann et al., 2003) and was chosen as a control to compare with previous studies using these same patient autoantibodies (Kreye et al., 2016; Michalski et al., 2024).

Cultures were treated overnight with patient antibodies at a concentration of 15 µg/mL, as in Kreye et al. (2016), and then imaged in 0 mM Mg^2+^ ACSF with TTX. In maximum minus minimum projection images (Fig. 8A) and in synaptic calcium traces (Fig. 8B), two antibodies, 003-102 and 008-218, decreased synaptic calcium transients compared to MGO-53 negative controls. The frequency of activity in cultures treated with antibodies 003-102 and 008-218 and number of active synapses also appeared decreased compared to MGO-53 controls and antibody 007-124, in raster plots (Fig. 8C). Antibody 007-124 did not change the number of active synapses compared to MGO-53 controls (p > 0.99; Fig. 8D), in line with its lower affinity compared to 003-102 and 008-218 (Ly et al. 2018). The number of active synapses trended toward a decrease after treatment with antibody 003-102 (p = 0.066; exclusion of one outlier resulted in p = 0.034) and was significantly decreased by antibody 008-218 (p < 0.0001). The finding that antibody 008-218 decreases the number of active synapses more than 003-102 is surprising, given that antibody 003-102 has a similar binding affinity for NMDARs compared to antibody 008-218 (Ly et al., 2018).

Patient antibodies 003-102 and 008-218 decreased the frequency of synaptic calcium transients at individual synapses compared to MGO-53 control treatments, while antibody 007-124 did not (Fig. 8E, top). The frequency of MGO-53-treated controls was 0.048 Hz. Antibody 003-102 decreased frequency to 0.039 Hz, by 21% compared to controls (0.79, 95% CI 0.70–0.90), and antibody 008-218 decreased frequency to 0.037 Hz, by 23% (0.77, 95% CI 0.68–0.87). Bootstrap resampling confirmed mixed-effects model results: both 003-102 and 008-218 antibodies decreased the frequency of synaptic calcium transients compared to MGO-53, while 007-124 showed no change (Fig. 8E, top; grey). The amplitude of synaptic calcium transients in control conditions was estimated to be 0.65 ΔF/F_0_. Antibodies 003-102 and 008-218 both decreased amplitude to 0.60 ΔF/F_0_ (003-102: 0.93, 95% CI 0.86–0.99; 008-218: 0.92, 95% CI 0.86–0.99). Antibody 007-124 did not affect amplitude (similar to its lack of effect on active synapse number and frequency, which were also unchanged).

## Discussion

Our automated high-throughput synaptic calcium imaging pipeline reports multiple parameters of synapse function, including the number of functional synapses and pre and postsynaptic function, for thousands of individual synapses. Using this pipeline, we collected data from 1,198,748 spontaneously active synapses and detected 10,228,773 unique synaptic calcium transient events without any manual ROI selection. We were able to attain such a dataset due to optimized automated analysis, in addition to semi-automated imaging, allowing multiple regions to be imaged in sequence on glass-bottom 24-well plates, which could be extended to 96- and 384-well plate formats. To accommodate such large datasets, we analyzed data using generalized linear mixed-effects models. These models were consistently comparable to stratified, clustered bootstrap resampling with replacement, indicating that both statistical tests give comparable results when analyzing average amplitude and frequency of synaptic calcium transient events. Examination of mixed-effects models supported modeling of the imaged region as a random effect; however, most of the variability in the data existed between synapses within the same field of view, rather than between regions.

We found that compounds that modulate postsynaptic receptors can also alter synaptic calcium transient frequency, but they simultaneously alter synapse number. For example, acute addition of glycine, a co-agonist of NMDARs, increased both frequency and synapse number, while acute addition of NMDAR antagonists such as ketamine, memantine, or APV decreased both frequency and number of active synapses. To differentiate whether changes in frequency are due to presynaptic or postsynaptic changes, reporting total active synapse numbers is crucial. All direct NMDAR agonists and antagonists changed the number of active synapses drastically within the same field of view compared to baseline recordings (p < 0.0001), whereas presynaptic modulators such as PDBu did not change the number of active synapses.

The blockade of synaptic calcium transients in the presence of APV verifies that they are NMDAR-dependent and aligns with results from previous studies (Andreae & Burrone, 2015; Metzbower et al., 2019; Prikhodko et al., 2024; Sinnen et al., 2016; Walker et al., 2017). The involvement of AMPARs in synaptic calcium transients is less clear. The AMPAR antagonist NBQX did not significantly change the frequency of calcium transients at individual synapses. However, there was a trend toward an increase in the number of active synapses (p = 0.088). Surprisingly, we observed an increase in amplitude of synaptic calcium transients in the presence of NBQX, contradicting results from previous studies (Walker et al., 2017; Metzbower et al., 2019, mentioned in text but data not shown). The observed increase in calcium transient amplitude could be attributed to the larger sample size investigated here and the use of automated ROI detection rather than manual detection. Biologically, an increase in amplitude caused by NBQX reflects well-established evidence that a single quanta does not saturate postsynaptic AMPARs and NMDARs (Mainen et al., 1999; McAllister & Stevens, 2000); therefore, with AMPARs blocked and more glutamate available to bind to NMDARs, an increase in amplitude of synaptic calcium transients is not unexpected.

The dendritic calcium transient events that were detected but not analyzed in most of the present study spread outward unidirectionally or bidirectionally along neuronal processes. Variability in spontaneous synaptic calcium transient propagation in dendrites often reflects differences in synaptic spine head and neck geometry, with larger spines more likely to propagate calcium into the dendritic arbor (Noguchi et al., 2005). It is also possible that dendritic calcium transients reflect immature synaptic sites. In DIV 4–5 hippocampal cultures, spontaneous calcium transients described and shown appear more elongated (Andreae & Burrone, 2015) than the synaptic puncta we primarily observe in our ‘mature’ cultures. In the presence of APV, ∼14% of synaptic event sites remained active, whereas ∼18% of dendritic event sites remained active. This result indicates that NMDARs are crucial for both event subtypes. However, calcium from non-NMDAR sources likely contributes to dendritic events, as evidenced by the fact that these dendritic events responded to APV differently compared to synaptic events: dendritic events maintained similar frequencies to control conditions, but displayed decreased amplitudes compared to controls and PDBu-treated conditions, while synaptic events had decreased frequency compared to controls and no change in amplitude compared to PDBu-treated conditions. Therefore, although NMDARs play a central role in the appearance of most dendritic events and the magnitude of transients, other calcium sources likely also contribute to their appearance as well. Previous studies have exclusively performed synaptic calcium imaging with a cocktail of antagonists, including NBQX, ryanodine, thapsigargin, and nifedipine to block non-NMDAR calcium entry to the dendritic arbor and report strictly NMDAR-dependent calcium transient events (Metzbower et al., 2019; Dean et al., 2022). It could be interesting to test the effects of blocking ryanodine receptors, store-operated calcium entry, or L-type voltage-gated calcium channels individually, rather than all of them indiscriminately, in future studies.

The differences in the effects of NMDAR autoantibodies we observe highlight the increased efficiency and sensitivity of our synaptic calcium imaging pipeline compared to other methods. The first paper characterizing the NMDAR patient antibody 003-102, for example, used a combination of immunocytochemistry to see decreases in NR1-positive clusters, electrophysiology to see decreases in NMDAR-dependent currents, and bath-applied NMDA with somatic calcium imaging to see decreased calcium influx (Kreye et al., 2016). Here, with a single experiment, we can reproduce these findings and extend readouts to synapse function, since we detected fewer active synapses and decreased amplitude of synaptic calcium transients at individual synapses following overnight application of patient antibodies. Changes in amplitude had previously been shown using whole-cell electrophysiology (Kreye et al., 2016; Michalski et al., 2024), which lacks single synapse resolution. In addition, we also found that two patient antibodies (003-102 and 008-218) decreased the frequency of spontaneous calcium transients, a novel finding. Ly et al. (2018) characterized the binding affinities of the NMDAR autoantibodies we tested and found 003-102 and 008-218 had similar binding affinities (with 003-102 formally higher), and 007-124 had the lowest affinity of the three. We found that overnight treatment with antibody 008-218 led to the largest decrease in the number of active synapses, even though it has a similar affinity for NMDARs as 003-102. This could be due to different mechanisms of action. Antibody 003-102 causes NMDAR internalization, whereas 008-218 causes NMDARs to adopt ‘non-active’ conformations where ion channels remain closed (Michalski et al., 2024). This could explain the larger decrease in number of active synapses observed in the presence of 008-218 compared to 003-102. These changes would not be observed as a change in immunostaining using fixed tissue, highlighting the advantage of our assay in testing functional effects of NMDAR autoantibodies on synapses.

Another NMDAR encephalitis antibody has been screened previously with synaptic calcium imaging, although no effect was observed (Dean et al., 2022). It is possible that the treatment time (10 minutes) was too short, since changes in spontaneous NMDAR current amplitude caused by other NMDAR antibodies occurred only after a 30-minute incubation in electrophysiological experiments (Michalski et al., 2024). Dean et al. also used only 1 μg/mL of the antibody, whereas our study and others tested concentrations at least an order of magnitude higher (Kreye et al., 2016; Michalski et al., 2024).

Several research areas could benefit immediately from such a functional synapse assay. NMDAR encephalitis is one such clinically relevant example for which our assay has already shown results. NMDAR encephalitis has an incidence of 1–5 per million people per year (Dalmau et al., 2019); however, it is likely underdiagnosed, with symptoms mimicking other classical psychiatric illnesses (Kayser & Dalmau, 2011). Diagnosis is currently only possible with an abnormal EEG, a spinal tap, and CSF screening with an antibody panel. We observed different effects of the three antibodies we tested, which could potentially underlie three distinct phenotypes of the disease. To better understand and characterize NMDAR encephalitis pathology, a functional antibody screen is essential. Our pipeline could be applied as well to screen patient CSF samples directly, potentially bypassing the need to purify autoantibodies, to directly report if patients show any NMDAR-specific pathologies and thus qualify for antibody-depleting therapies. Our automatic analysis not only avoids manual intervention and potential experimenter selection bias but also allows the detection of more nuanced changes due to the large, unrestricted sample size analyzed.

Our pipeline could also be used to study several conditions known to affect synaptic activity, such as schizophrenia (Glantz & Lewis, 2000; Park et al., 2022), autism spectrum disorder (Monteiro & Feng, 2017), Parkinson’s disease (Reyes-Resina et al., 2024), or Alzheimer’s Disease (Escamilla et al., 2024; Prikhodko et al., 2024). Additionally, this pipeline could be used to investigate how synaptic calcium transients differ between neuronal subtypes, for example, by using different GCaMP6f constructs driven by the CaMKII promoter (for excitatory neurons) or the Dlx promoter (for inhibitory neurons). Sparse transduction could also be used to limit neurite overlap (Metzbower et al., 2019), which would allow synaptic activity from individual neurons to be studied and potential differences between distal and proximal synapses to be observed before and after compound treatments at scale (Walker et al., 2017). GCaMP6f could also be co-expressed with a synapse-targeted fluorophore of a different color to quantify the proportion of total synapses that are active and whether extrasynaptic NMDARs play a role in the calcium transients observed.

Our primary goal was to analyze synaptic calcium transients in an automated fashion. With this pipeline, we streamline the analysis of the activity of individual synapses and routinely visualize in a single video what other investigators have used for their entire datasets. In doing so, we open the possibility to study individual synapses on a scale not previously realized. High-throughput large-scale analyses of synapse function could offer new and interesting insights into plasticity and neural network formation in vitro and in vivo, as well as improve characterization of disease models and development of compounds that prevent or reverse synaptic dysfunction.

## Author Contributions

John Carl Begley performed all calcium imaging experiments, created all Python, R, ImageJ, and CellProfiler pipelines, and optimized all for automation. Paul Turko provided cell cultures for experiments and assisted with experiment design. Harald Prüss provided encephalitis patient-derived NMDAR auto-antibodies. Camin Dean and Paul Turko co-supervised the project. The manuscript was written by John Carl Begley with assistance from Camin Dean and was edited by Paul Turko and Harald Prüss.

## Acknowledgements

We thank the Shared Primary Neuron Facility of the Institute for Integrative Neuroanatomy at the Charité and their technicians, Birgit Metze, Petra Loge, and Marion Möbes, for providing the cell cultures for all experiments. We thank Justus Donner, Fabian Ottolin, Marti Ritter, Hatem Oraby, and Hung Lo for their technical assistance with improving the Python code and automation of the pipelines. We thank Kerstin Rubarth for her assistance in choosing statistical tests for our datasets. We thank the Advanced Medical BIOimaging Core Facility of the Charité-Universitätsmedizin Berlin (AMBIO) for support in the acquisition of the imaging data.

---

Group sizes supplemental table.

| Experiment | Cultures | Wells Imaged | Regions Imaged | Synaptic Events | Dendritic Events | Total Events |
| --- | --- | --- | --- | --- | --- | --- |
| Ctrl →<br>PDBu →<br>APV | 3 | 9 | 27 | 61,245<br>64,757<br>8,606 | 6,479<br>9,393<br>1,186 | Ctrl: 67,855<br>PDBu: 74,150<br>APV: 9,973 |
| Ctrl →<br>Glycine | 3 | 9 | 27 | 67,587<br>76,467 | 6,292<br>8,205 | Ctrl: 70,907<br>Glycine: 83,465 |
| Ctrl →<br>NBQX | 3 | 9 | 27 | 62,934<br>64,831 | 5,107<br>6,912 | Ctrl: 68,680<br>NBQX: 71,743 |
| Overnight MGO-53 control<br>antibody | 4 | 15 | 31 | 49,322 | 6,947 | 56,512 |
| Overnight anti-NMDAR<br>antibody 003-102 | 4 | 15 | 31 | 37,431 | 3,833 | 41,512 |
| Overnight anti-NMDAR<br>antibody 007-124 | 4 | 15 | 31 | 50,736 | 7,004 | 58,213 |
| Overnight anti-NMDAR<br>antibody 008-218 | 4 | 15 | 31 | 20,298 | 3,367 | 24,605 |
| Ctrl →<br>Ketamine [1 μM] | 3 | 9 | 27 | 58,841<br>47,759 | 5,543<br>2,955 | 64,543<br>50,912 |
| Ctrl →<br>Memantine [1.5 μM] | 3 | 9 | 27 | 64,161<br>37,718 | 6,515<br>1,996 | 70,820<br>40,904 |
| Ctrl →<br>DMSO veh. | 3 | 9 | 27 | 48,605<br>50,449 | 4,933<br>4,756 | 55,538<br>55,470 |
| Overnight DMSO | 3 | 4 | 15 | 11,227 | 1,126 | 12,353 |
| Overnight Ketamine [1 μM] | 5 | 10 | 38 | 37,958 | 2,698 | 42,476 |
| Overnight Ketamine [10 μM] | 3 | 6 | 24 | 15,804 | 848 | 16,729 |
| Overnight Memantine [1.5 μM] | 4 | 8 | 29 | 33,806 | 2,406 | 36,252 |
| Overnight Memantine [10 μM] | 3 | 6 | 23 | 19,299 | 1,600 | 21,533 |

